# Inhibitory effects of dopamine agonists on pain-responsive neurons in the central nucleus of the amygdala

**DOI:** 10.1101/2025.03.10.642168

**Authors:** Robert J. Heuermann, Robert W. Gereau

## Abstract

The central nucleus of the amygdala (CeA) is a heterogenous region of primarily GABAergic neurons that contributes to numerous behaviors, including fear learning, feeding, reward, and pain. Dopaminergic inputs to the CeA have been shown to regulate many of these behaviors, but how dopamine exerts these effects at the cellular level has not been well characterized. We used the Targeted Recombination in Active Populations (TRAP) mouse line to fluorescently label pain-responsive CeA neurons, and then targeted these cells for patch-clamp recordings in acute slices to test the effects of dopamine agonists. The D1 agonist SKF-38393 and D2 agonist quinpirole both had inhibitory effects, reducing the input resistance and evoked firing and increasing rheobase of labeled CeA neurons. Both agents also inhibited the NMDA component of excitatory postsynaptic currents (EPSCs) evoked by basolateral amygdala (BLA) stimulation, but did not affect the AMPA component. D1 activation, but not D2, also appeared to have a presynaptic effect, increasing the frequency of spontaneous EPSCs. These results provide new insights into how dopamine regulates activity within pain-responsive CeA networks.

**NEW & NOTEWORTHY:** Dopamine is known to regulate activity within the central amygdala (CeA), an important region for central pain processing. However, its effects at the cellular level have not been well characterized. We targeted pain-responsive CeA neurons for patch-clamp recordings to examine the cellular and synaptic effects of D1 and D2 agonists. Activation of either D1 or D2 receptors induced inhibitory effects, suggesting dopamine signaling in CeA dampens pain-related activity and could be a target for analgesics.

## INTRODUCTION

Pain is a complex phenomenon involving both peripheral sensory input and central affective processing. Inadequate pain management represents one of the greatest unmet clinical needs contributing to patient suffering and healthcare costs worldwide (Zimmer et al. 2022). The affective component of pain in particular is not as well understood as the sensory pathways, yet is often the primary driver of morbidity and the development of chronic pain disorders (Lindsay et al. 2021). A key node within the affective pain circuitry is the central nucleus of the amygdala (CeA) (Neugebauer et al. 2020). The CeA consists of primarily GABAergic neurons, similar to the striatum, but with more heterogeneous molecular and electrophysiological phenotypes (Adke et al. 2020; Kim et al. 2017; Li et al. 2022; Wang et al. 2023). CeA neurons are highly interconnected in both local circuits and with many other cortical and subcortical areas (Zhang et al. 2021). The CeA is thus poised to integrate direct nociceptive inputs from the parabrachial nucleus with higher-order information about emotional state, perceived threats, and internal cues like hunger and thirst (Moscarello and Penzo 2022).

Like the striatum, the CeA receives midbrain dopaminergic inputs from the substantia nigra, ventral tegmental area, and periaqueductal gray (Casey et al. 2023; Poulin et al. 2018; Zhao et al. 2022). While dopamine has received increasing attention as a modulator of pain signaling at various levels – including within the spinal cord (Barasi et al. 1987; Taniguchi et al. 2011), striatum (Magnusson and Fisher 2000), nucleus accumbens (Ren et al. 2020), and cortex (Huang et al. 2020; Lançon et al. 2021) — there has been minimal investigation into its potential role in pain processing within the CeA (Huang et al. 2022). Dopamine has been shown to influence other functions of the CeA, such as anxiety-like (László et al. 2020; Liu et al. 2011; Mora et al. 2005) and appetitive (Jin et al. 2020) behaviors, fear learning (Bundel et al. 2016; Groessl et al. 2018; Jo et al. 2018), and attention (Lee et al. 2006; Smith et al. 2015). Several studies have also identified expression of both D1 and D2 receptors in CeA neurons (Casey et al. 2022; Kim et al. 2017; Lu et al. 2021). However, the response to dopamine in the CeA has not been well characterized at the cellular level.

Here, we sought to characterize the cellular and synaptic effects of D1 and D2 agonists on CeA neurons. We specifically targeted putative pain-responsive neurons using the Targeted Recombination in Active Populations (TRAP) mouse line (Guenthner et al. 2013). In patch-clamp recordings from acute CeA slices, we found that both D1 and D2 agonists produced inhibitory effects on intrinsic neuronal properties of pain-responsive neurons. Both agonists also inhibited the NMDA component of stimulated excitatory inputs from the basolateral amygdala (BLA) but had no effect on the AMPA component. These findings enhance our understanding of dopamine’s role in the CeA, particularly in the context of central pain processing.

## MATERIALS AND METHODS

### Animals

TRAP2 (Fos^tm2.1(icre/ERT2)Luo^/J) and Ai14 mouse lines were sourced from Jackson labs and maintained locally. All mice in this study were F1 offspring of crosses of homozygotes from these two lines. All experiments were conducted in accordance with National Institute of Health guidelines and with approval from the Animal Care and Use Committee of Washington University School of Medicine. Both male and female mice were used equally.

### Targeted Recombination in Active Populations (TRAP)

Mice aged 6-10 weeks were separated from their littermates and singly housed for 7-10 days to minimize confounding stimuli during the TRAP procedure. Both male and female mice were used. Mice were acclimated to handling for 5-7 days prior to the induction procedure, and at least 3 consecutive days before induction. Acclimation included applying gentle pressure to each hindpaw and being scruffed and touched on the abdomen to simulate intraperitoneal (i.p.) injection. The day of induction, a hemostat was used to apply noxious pinches to each hindpaw (2 pinches per paw of 5 s each). Animals displayed clear nocifensive behaviors of freezing, vocalizations, and/or biting the hemostat. Mice were then returned to their home cage, and 30-60 minutes later 4-hydroxytamoxifen (Sigma-Aldrich or HelloBio) dissolved in corn oil (5 mg/mL) was administered (20 mg/kg i.p.).

### Tissue clearing

3-42 days after the TRAP procedure, mice were deeply anesthetized with isoflurane, transcardially perfused with 20 mL of chilled PBS, then decapitated and brains rapidly dissected into chilled PBS. Coronal slices (300-500 µm) were obtained using a Compresstome VF-210-0Z (Precisionary Instruments). Slices were fixed overnight at 4ºC in 4% paraformaldehyde in PBS, and then stored at 4ºC in PBS until further processing. Slices were then cleared using the ScaleSQ method (Hama et al. 2015). Briefly, PBS was removed and slices were immersed in ScaleSQ solution (9.1M urea, 22.5% sorbitol, 5% Triton X-100) for 2-3h rocking at 37ºC. ScaleSQ was then removed and ScaleS4 added (4M urea, 40% sorbitol, 10% glycerol, 15% dimethylsulfoxide) for 2-3h rocking at room temp. Slices were left in ScaleS4 and stored at 4ºC until mounting and imaging. ScaleS4 was used as the mounting medium. Slices were mounted on glass slides and covered with glass coverslips. Slices were noted to rapidly re-opacify if exposed to nail polish during mounting, so the coverslip edges were first coated in ScaleS4 with 3% low-melt agarose (melted at 90ºC and applied with a paintbrush) prior to sealing with clear nail polish.

### Imaging

Epifluorescence images of TdTomato-labeled cells were obtained on a Leica DM6b system with LAS-X software (v3.7). Images were first obtained as Z-stack (50 µm step size) tile scans of each slice, auto-stitched in LAS-X, and converted to maximum-intensity projections for further analysis. This was able to clearly resolve all cells within each 300-500 µm section. Atlas registration and semi-automated regional cell counting was performed using the QUINT workflow (Yates et al. 2019), after training the Ilastik object classifier on sample images. Several slices for each sample were manually reviewed to ensure accuracy. Cell counts were analyzed in Excel (Microsoft 365). For normalized cell counts in Figure 1C, the total number of TdTomato-labeled cells from all slices containing CeA was used as the denominator for each sample. The experimenter was blinded to the condition of each animal until after all images and cell counts were acquired.

**Figure 1.**
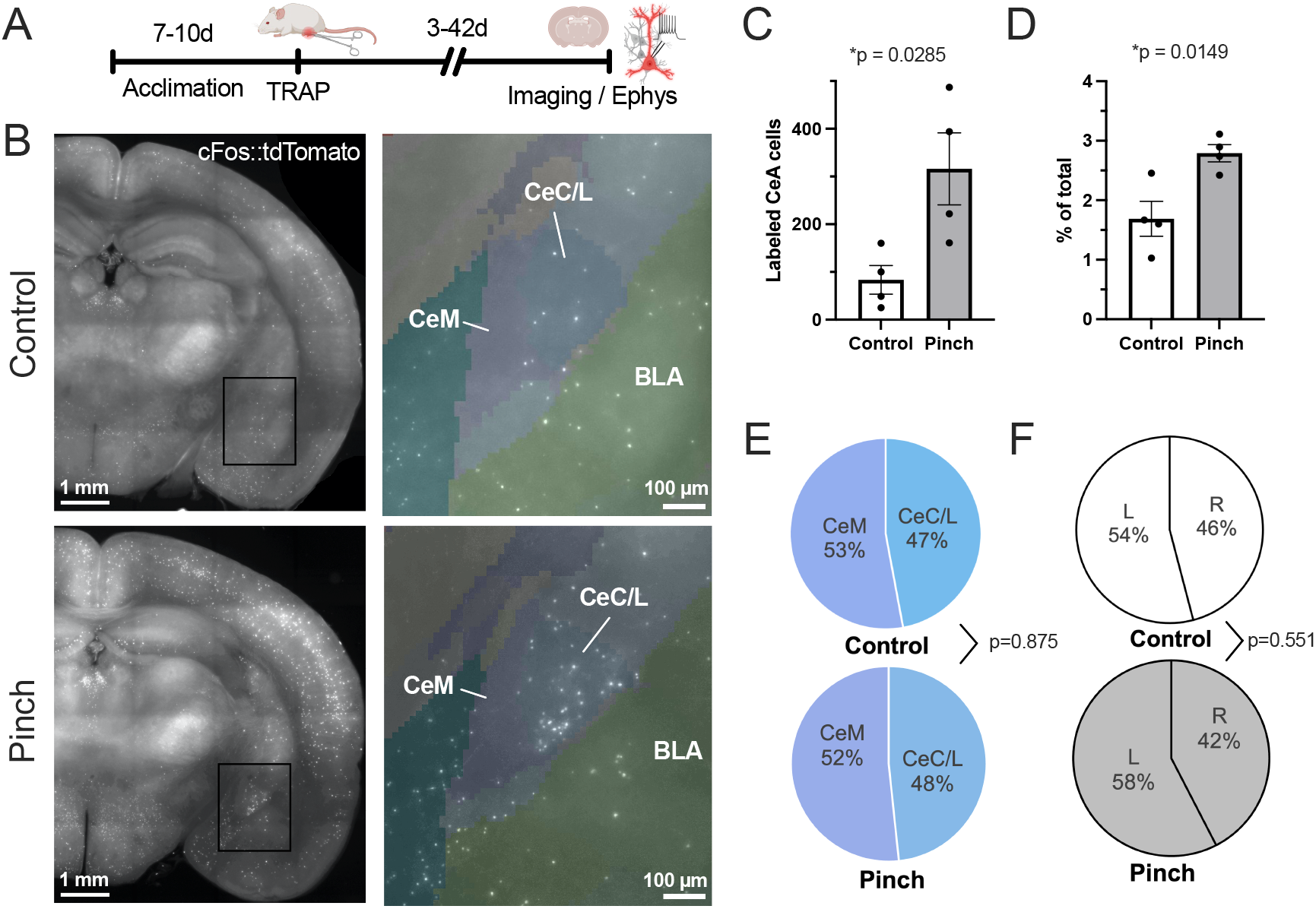
Targeted Recombination in Active Populations (TRAP) to label pain-responsive cells in the CeA. **(A)** Schematic of timeline for TRAP procedure. **(B)** Representative images of tdTomato labeling in coronal slices from control mice (top) and mice that received a painful paw pinch (bottom). Left: 5x tile-scan images. Right: Magnified view of the boxed regions, with color overlay of the regional segmentations generated by the QUINT workflow. CeM: medial division of CeA; CeC/L: capsular/lateral division of CeA; BLA: basolateral amygdala. **(C**,**D)** Absolute (B) and normalized (C) counts of tdTomato+ cells in control vs. pinch mice (n=4; unpaired t-test) **(E**,**F)** Control and pinch conditions did not differ in the distribution of labeled cells (D) within CeA subdivisions (n=4; unpaired t-test of % CeC/L values, p=0.875); or (E) between right and left CeA (n=4; unpaired t-test of % Left values, p=0.551).

### Electrophysiology

3-42 days after the TRAP procedure, mice were deeply anesthetized with ketamine / xylazine / acepromazine (100 / 8 / 2.5 mg/kg i.p.) then transcardially perfused with 20 mL of chilled N-methyl-D-glucamine (NMDG)-based slicing solution containing (in mM): 93 NMDG, 2.5 KCl, 1.25 NaH2PO_4_, 30 NaHCO_3_, 20 HEPES, 25 glucose, 5 ascorbic acid, 2 thiourea, 3 sodium pyruvate, 5 MgSO_4_, 0.5 CaCl_2_, 12 N-acetylcysteine; with pH adjusted to 7.3 using HCl, and continuously bubbled with carbogen (95% O_2_ / 5% CO_2_). Brains were dissected and mounted for making Compresstome slices (300 µm), then transferred to a recovery chamber with NMDG solution warmed to 32ºC for 10-12 min, and finally transferred to 32ºC artificial cerebrospinal fluid (ACSF) containing (in mM): 124 NaCl, 2.5 KCl, 1.25 NaH_2_PO_4_, 24 NaHCO_3_, 5 HEPES, 12.5 Glucose, 1 MgCl_2_, 2 CaCl_2_; with pH adjusted to 7.3 using NaOH, and continuously bubbled with carbogen. The holding chamber was allowed to gradually equilibrate to room temp for 1 hour prior to recordings.

Slices were then transferred to a recording chamber perfused continuously with ACSF (2.0 mL/min, continuously bubbled with carbogen). Recordings were performed at room temperature. CeA neurons were visualized on an Olympus BX50WI microscope equipped with differential interference contrast optics and epifluorescence filters. Thin-walled borosilicate pipettes (3-6 MΩ) were pulled on a Sutter Instruments P-97. For current-clamp recordings, internal solution contained (in mM): 120 K-gluconate, 5 NaCl, 2MgCl_2_, 0.1 CaCl_2_, 10 HEPES, 1.1 EGTA, 4 Na_2_ATP, 0.4 Na_2_GTP, 15 Na_2_Phosphocreatine, and 0.1% biocytin; adjusted to pH 7.3 with KOH; 295 mOsm). For synaptic recordings, internal solution contained: 110 D-gluconic acid, 110 CsOH, 0.1 CaCl_2_, 10 HEPES, 1.1 EGTA, 4 MgATP, 0.4 Na_2_GTP, 10

Na_2_Phosphocreatine, 8 tetraethylammonium-Cl, 3 QX314-Br, and 0.1% biocytin; adjusted pH to 7.3 with HCl; 290 mOsm). A bipolar stimulating electrode (100 MΩ, World Precision Instruments) was placed in the basolateral amygdala (BLA) 100-200 µm from the recorded neuron using a manual micromanipulator. Biphasic stimulation (100-500 µA, 1 ms pulse width) was delivered using an A385 stimulus isolator unit (World Precision Instruments). Picrotoxin (100 µM) was included in the ACSF to block GABA_A_ currents during synaptic but not current-clamp recordings.

Recordings were low-pass filtered at 3 kHz and sampled at 10 kHz with a MultiClamp 700B amplifier, Digidata 1440 digitizer, and pClamp 11 software (Molecular Devices. Pipette capacitance was compensated prior to breaking into each cell. Series resistance (Rs) was monitored throughout each recording, and cells discarded if it exceeded 40 MΩ. Bridge balance was compensated in current clamp recordings. Series resistance was not compensated in voltage clamp. Membrane potential values were not corrected for a calculated liquid junction potential of 14 mV. In current clamp, after recording the initial resting membrane potential, a holding current was added to achieve a baseline potential of -70 ± 1 mV. A series of current steps from -200 to +200 pA (1 s duration, 25 pA step size) was then delivered for measurement of input resistance (Rin) and evoked action potentials (APs). For measurement of rheobase (the minimum current required to elicit an AP), a linear ramp of depolarizing current was applied over 3 s. The ramp peak was initially adjusted for each cell such that the first AP occurred approximately halfway through the sweep, and kept constant thereafter. For synaptic recordings, the stimulus intensity was initially adjusted for each cell to achieve evoked excitatory post-synaptic potentials (EPSCs) of 100-400 pA (at -70 mV holding potential), and kept constant thereafter. The AMPA component of the EPSC was measured as the peak value at -70 mV. The NMDA component was estimated as the amplitude 50 ms after the stimulus during sweeps at +40 mV holding potential. Paired-pulse ratio (PPR) was obtained at -70 mV from 2 stimuli at 20 Hz.

After obtaining baseline recordings for each cell, the bath perfusion source was switched to ACSF containing quinpirole (10 µM) or SKF-38393 (10 nM). 5 minutes was allowed for drug wash-on, and then the recording protocol was repeated. All chemicals were from Sigma-Aldrich unless otherwise stated.

### Data analysis

Recording data was imported into Igor Pro 8 using custom routines adapted from the NeuroMatic plugin (Rothman and Silver 2018), and further analyzed with custom functions. R_in_ was calculated as the slope of the linear regression fit of the steady-state voltage change for steps of -50, - 25, and 0 pA. More negative steps were not included for R_in_ calculations due to prominent inward rectification in some CeA neurons. Rheobase was derived from the ramp protocol above as the instantaneous current when the first AP reached threshold. The first peak of the second derivative of the AP waveform was used to define threshold. The rheobase from 3 consecutive sweeps was averaged for each cell. For evoked synaptic recordings, 5 consecutive sweeps were averaged for each measurement (I_AMPA_, I_NMDA_, and PPR). Spontaneous EPSCs were detected by manual inspection of 2-minute recordings at a -70 mV holding potential after applying a 100 Hz low-pass filter. Cumulative histograms were generated from pooled events from all cells.

### Statistics

Statistical analyses were performed using Prism 10 (GraphPad). Specific tests are listed in the figure legends. Data are plotted as mean ± s.e.m.

## RESULTS

Due to the broad phenotypic heterogeneity within CeA neurons, we sought to selectively target cells involved in pain signaling. To accomplish this, we used the Targeted Recombination in Active Populations (TRAP) mouse line (Guenthner et al. 2013), which express tamoxifen-dependent Cre recombinase under control of the c-Fos promoter. Cre expression is thus up-regulated in response to neuronal activity but is inactive unless there is concurrent administration of tamoxifen or 4-hydroxytamoxifen (4-OHT). We crossed TRAP mice with the Ai14 reporter line, enabling permanent tdTomato labeling of neurons active during the induction protocol. Induction consisted of a noxious pinches delivered to the bilateral hindpaws using a hemostat. This protocol produced robust expression of tdTomato-labeled neurons in the CeA (**Figure 1A**), with significantly more labeled cells in the pinched group compared to no-pinch controls (which instead received gentle hindpaw pressure during induction) (**Figure 1B-C**). Labeled cells were roughly equally distributed between the medial (CeM) and capsular/lateral (CeC/L) subdivisions of the CeA, and this distribution did not differ between the pinch and no-pinch conditions (**Figure 1D**; control: 47.1 ± 7.3% CeC/L; pinch: 48.3 ± 2.3%). We also did not observe any difference between the right and left CeA in either condition (**Figure 1E**; control: 45.9 ± 4.0% right; pinch: 42.4 ± 3.8%). This is noteworthy because some aspects of pain signaling within the CeA exhibit hemispheric lateralization (Allen et al. 2022; Carrasquillo and Gereau 2008; Ji and Neugebauer 2009).

We next targeted pinch-labeled CeA cells for patch clamp recordings in acute slices to test the effects of dopamine agonists (**Figure 2A**). We specifically targeted the CeC/L subdivision as it has been most associated with nociceptive processing (Neugebauer et al. 2020). Prior studies suggest CeC/L neurons primarily express D2 receptors (Kim et al. 2017; Lu et al. 2021), so we first tested the D2 agonist quinpirole (10 µM). Compared to baseline recordings, wash-on of quinpirole decreased input resistance (**Figure 2D-E**). The number of action potentials (AP) evoked by depolarizing current steps was also reduced (**Figure 2F**), and rheobase (the minimum current needed to induce an AP) was increased (**Figure 2G-H**). There was no significant change in resting membrane potential (RMP, **Figure 2C**) or AP threshold (**Figure 2H**). Together, these data support an inhibitory effect of quinpirole on CeC/L neurons in a subpopulation enriched for pain-responsiveness.

**Figure 2.**
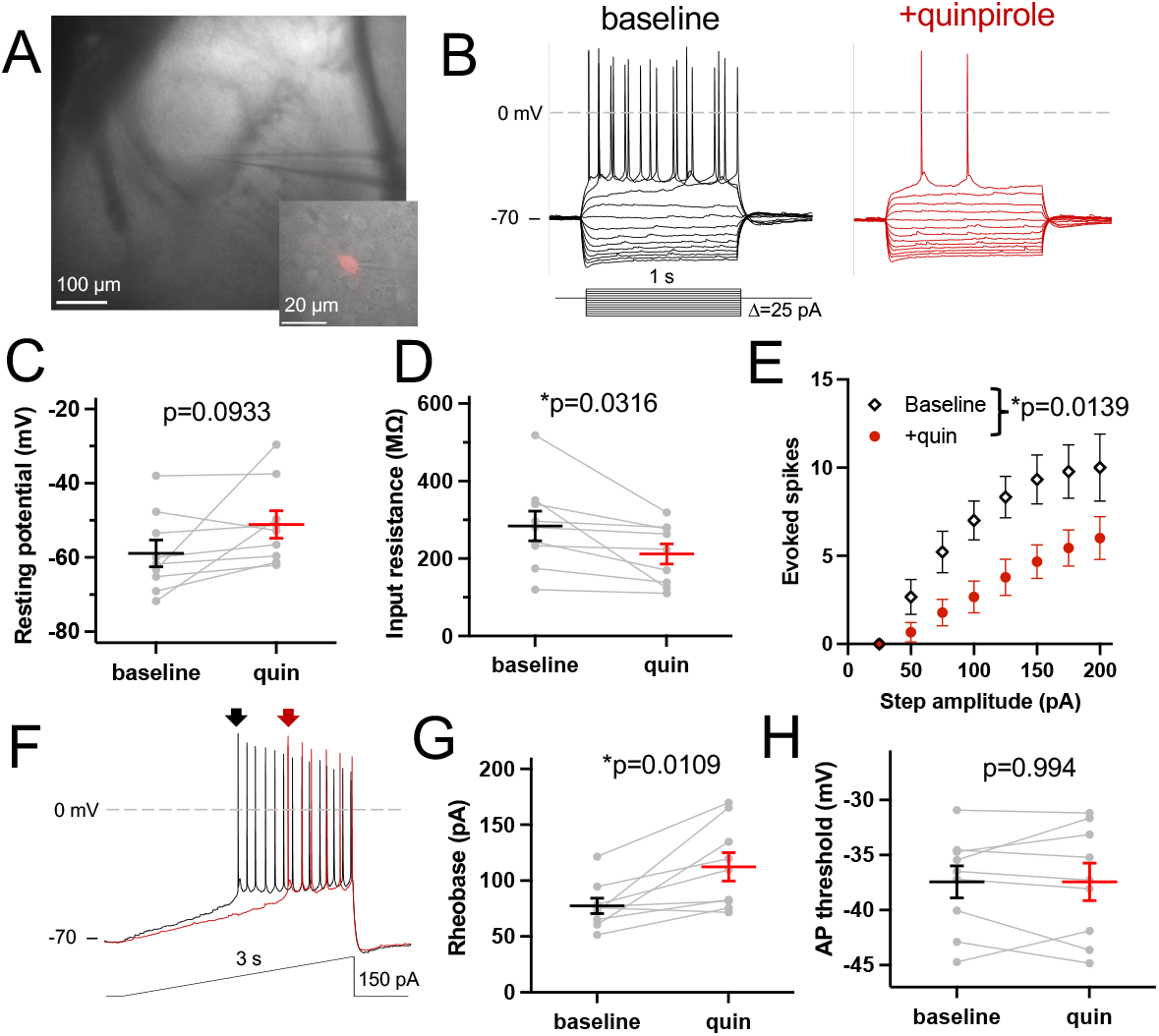
The D2 agonist quinpirole has inhibitory effects on pain-responsive CeA neurons. **(A)** Representative low-power (5x) image showing recording configuration for targeting CeA neurons. Inset: 40x brightfield image with red fluorescence overlay showing successful recording of a tdTomato^+^ neuron. **(B)** Representative traces of current step recordings before (black) and after (red) applying quinpirole (10 µM). **(C-E)** Group data for resting membrane potential (C), input resistance (D), and number of evoked spikes (E) of baseline vs. quinpirole recordings. **(F)** Representative traces of ramp protocol used for rheobase calculations. Arrows indicate first action potential, showing increased rheobase after quinpirole (red). **(G**,**H)** Group data for rheobase (G) and AP threshold (H). Paired t-test for C, D, G, and H (n=9). 2-way repeated-measures ANOVA for E (n=9; F(1,8)=9.83). Data presented as mean ± SEM.

Surprisingly, application of the D1 agonist SKF-38393 (10 nM) had largely similar effects (**Figure 3**). Here, SKF-38393 induced a significant depolarization in RMP (**Figure 3B**). Like quinpirole, it also reduced input resistance and evoked APs, increased rheobase, and had no effect on AP threshold (**Figure 3C-G**). Thus, D1 activation also has a net inhibitory effect on putative pain-responsive CeA neurons.

**Figure 3.**
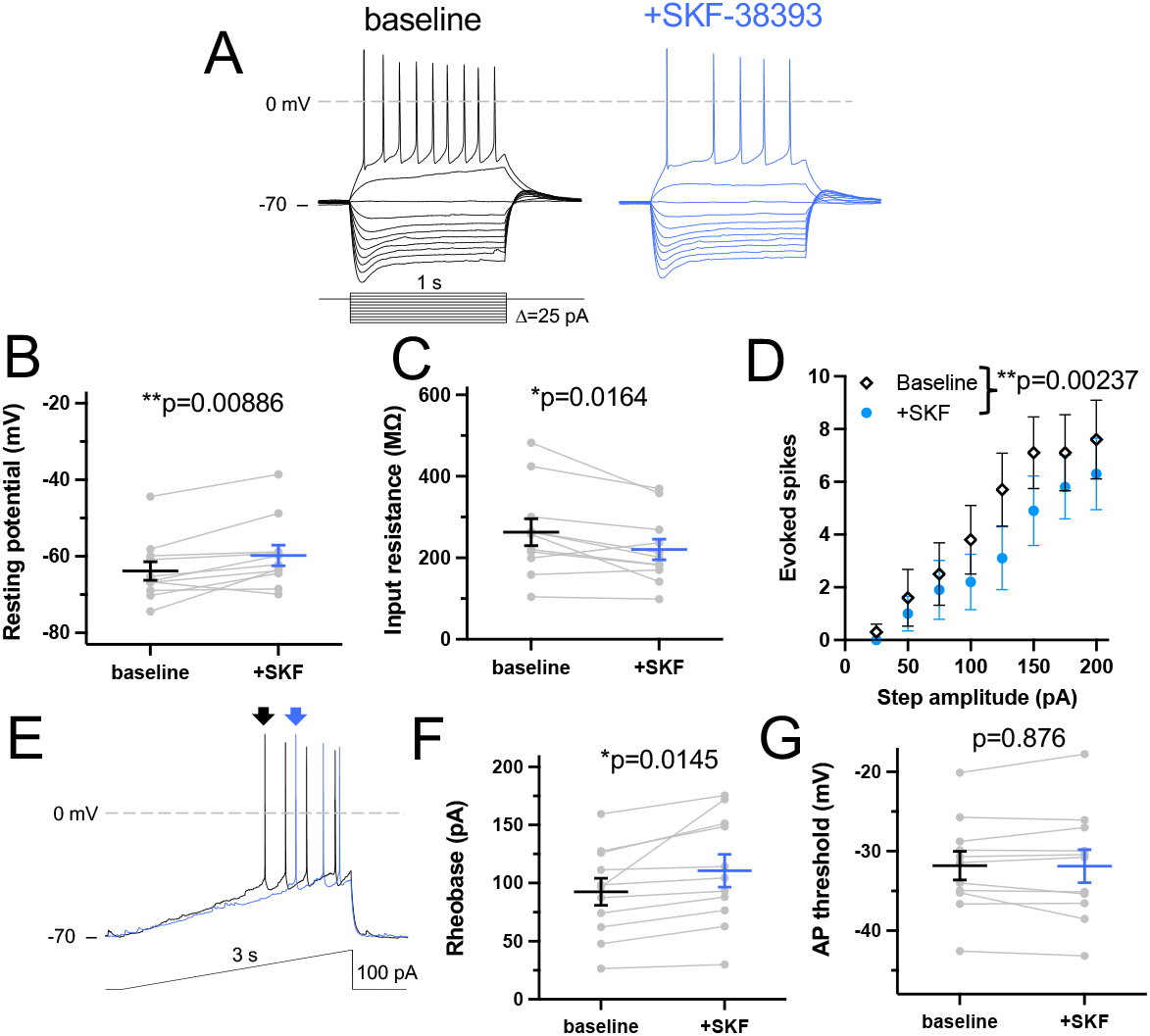
The D1 agonist SKF-38393 has inhibitory effects on pain-responsive CeA neurons. **(A)** Representative traces of current step recordings before (black) and after (blue) applying SKF-38393 (10 nM). **(B-D)** Group data for resting membrane potential (B), input resistance (C), and number of evoked spikes (D) of baseline vs. SKF recordings. **(F)** Representative traces of ramp protocol used for rheobase calculations. Arrows indicate first action potential, showing increased rheobase after SKF (blue) **(F**,**G)** Group data for rheobase (F) and AP threshold (G). Paired t-test for B, C, F, and G (n=11). 2-way repeated-measures ANOVA for D (n=11; F(1,10)=16.30). Data presented as mean ± SEM.

We also tested the effects of dopamine agonists on synaptic inputs from the basolateral amygdala (BLA), which supply one of the strongest inputs to CeA neurons. Electrical stimulation of the BLA during slice recordings produces robust evoked excitatory postsynaptic currents (EPSCs). Quinpirole application had no effect on EPSC amplitude at a holding potential of -70 mV but caused a significant reduction at +40 mV, reflecting the AMPA and NMDA components of the response, respectively (**Figure 4A-C**). There was a corresponding ∼3-fold increase in AMPA:NMDA ratio (**Figure 4D**). There was no effect on paired-pulse ratio (**Figure 4E**), suggesting this effect was postsynaptically mediated. There was also no difference in the frequency or amplitude of spontaneous EPSCs recorded at -70 mV(**Figure 4F-G**), further consistent with a postsynaptic mechanism.

**Figure 4.**
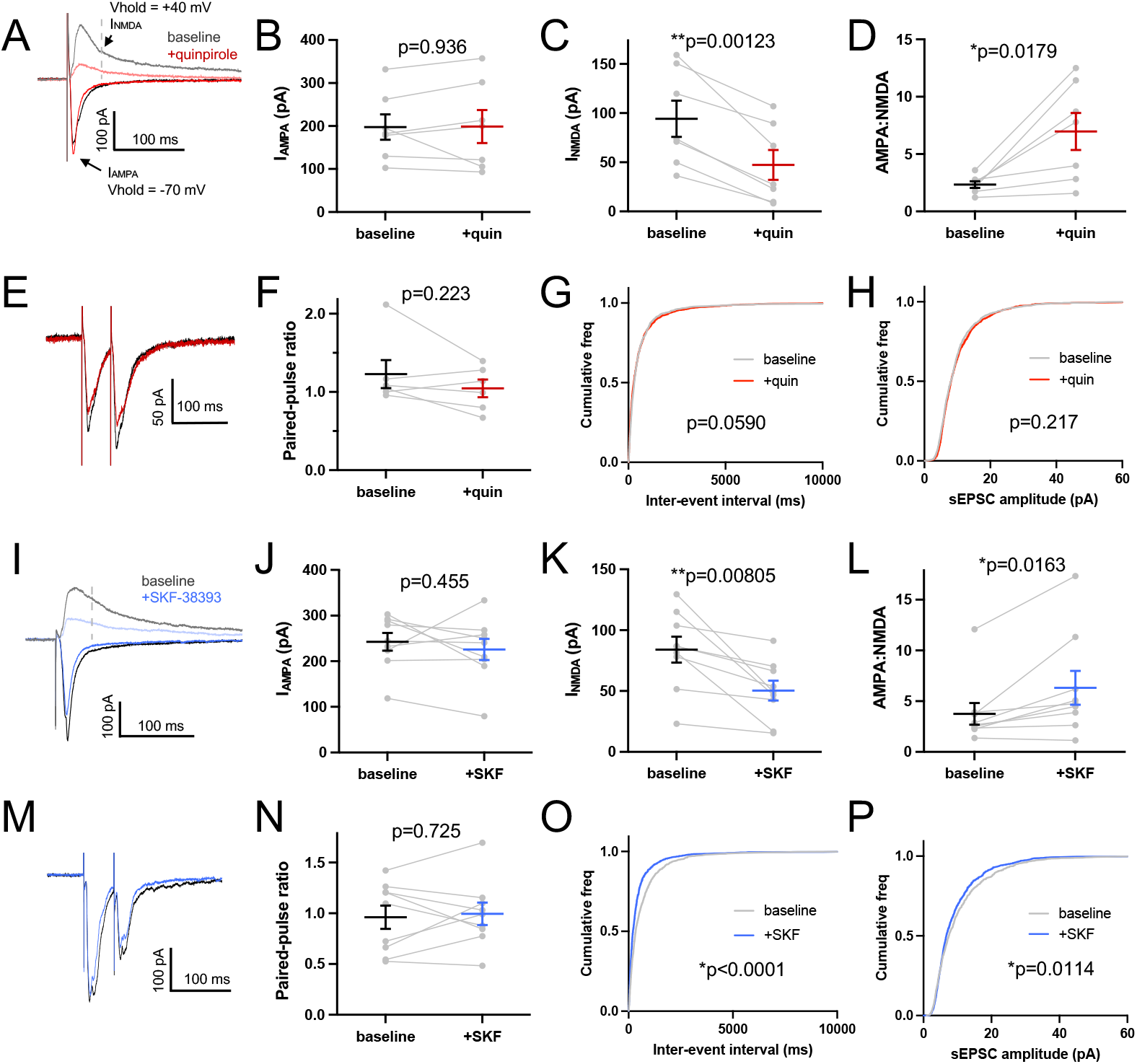
Effects of quinpirole and SKF-38393 on synaptic inputs from BLA. **(A)** Representative baseline-subtracted traces of evoked EPSCs at -70 mV (negative currents) and +40 mV (positive currents), before (gray) and after (red) applying quinpirole. **(B-D)** Group data for I_AMPA_, I_NMDA_, and AMPA:NMDA ratio with quinpirole (n=7; paired t-test). **(E**,**F)** Representative traces and group data for paired-pulse ratio before and after quinpirole (n=7; paired t-test). **(G**,**H)** Cumulative histograms of inter-event interval and amplitude of spontaneous EPSCs before and after quinpirole (n=7; Kolmogorov-Smirnov test). **(I)** Representative evoked EPSC traces before (gray) and after (blue) SKF-38393. **(J-L)** Group data for I_AMPA_, I_NMDA_, and AMPA:NMDA ratio with SKF (n=9; paired t-test). **(M**,**N)** Representative traces and group data for paired-pulse ratio before and after SKF (n=9; paired t-test). **(O**,**P)** Cumulative histograms of interevent interval and amplitude of spontaneous EPSCs before and after SKF (n=9; Kolmogorov-Smirnov test).

SKF-38393 had a similar effect on evoked EPSCs, with reduction in the NMDA component, no change in the AMPA component, and no change in paired-pulse ratio (**Figure 4H-N**). However, in this case there was also a significant increase in spontaneous EPSC frequency, as well as a slight decrease in amplitude (**Figure 4O-P**). D1 activation thus appears to have mixed effects on excitatory inputs to the CeA via both pre- and post-synaptic actions.

## DISCUSSION

In this study, we characterized the cellular effects of dopamine agonists on CeA neurons, targeting a population enriched for pain-responsive cells. The D1 agonist SKF-38393 and D2 agonist quinpirole both had net inhibitory effects on intrinsic excitability – reducing input resistance, increasing rheobase, and reducing evoked action potential firing. Both agents also reduced the NMDA component of excitatory inputs from the BLA, with no effect on the AMPA component. The effects of quinpirole appeared to be entirely post-synaptic, but SKF-38393 increased the frequency of spontaneous EPSCs, suggesting an additional presynaptic effect.

There is growing interest into how dopamine signaling affects the various functions of the CeA. D1 antagonists infused into the CeA reduce anxiety-like behavior (Fernandes et al. 2021; Liu et al. 2011; Mora et al. 2005), whereas D2 antagonists (László et al. 2020; Mora et al. 2016) or regional knockout of the D2 receptor in CeA (Casey et al. 2022) increase measures of anxiety. Similarly, D2 inhibition in the CeA caused generalization of conditioned fear (Bundel et al. 2016), and optogenetic activation of dopamine inputs from the VTA reduced generalization (Jo et al. 2018). D1 activation has also been implicated in reinforcing behaviors with food rewards (Jin et al. 2020) and models of addiction (Jokar et al. 2023; Kim and Lattal 2019). Finally, activating VTA terminals in the CeA was shown to reduce mechanical hypersensitivity in a model of post-incisional pain (Huang et al. 2022).

Despite this converging evidence that dopamine plays an important role in CeA function, there has been relatively limited investigation into the physiological basis for these effects. One study showed an increase in spontaneous EPSC frequency after dopamine application (Silberman and Winder 2013), and our results with SKF-38393 now suggest this is D1-mediated. Another study examining inhibitory currents found a D1-mediated reduction in evoked IPSC amplitude (Naylor et al. 2010), and exogenous dopamine also decreased the frequency of spontaneous IPSCs. Interestingly, dopamine did not affect the input resistance of CeA neurons in this study, in contrast to our results with both quinpirole and SKF-38393. This discrepancy could be due to the considerable heterogeneity among CeA neurons, such that our recordings from a pain-responsive population captured an effect that was not evident with a nonselective sampling. Along these lines, Groessl et al. (2018) found that dopamine augmented long-term potentiation (LTP) of BLA inputs to a population of CeA neurons that express somatostatin (SST), but not those expressing PKCδ. Dopamine was also shown to potentiate BLA inputs to CeA in slices from cocaine-withdrawn rats, but not saline controls (Krishnan et al. 2011). In both cases, a D1 antagonist blocked the potentiating effect of dopamine. This contrasts with our finding that both D1 and D2 agonists inhibit the NMDA component of BLA EPSCs, which could again be attributable to cell-type heterogeneity. However, together these results suggest the intriguing possibility that the effect of dopamine on CeA circuits is state-dependent – further augmenting synapses that have undergone recent potentiation (via either high-frequency stimulation or cocaine withdrawal in the above studies), but inhibiting plasticity at less active synapses. This would have the effect of increasing the signal-to-noise ratio of select pathways, and could explain the reported effects of CeA dopamine to enhance learned behaviors (László et al. 2020; Yang et al. 2023) and reduce fear generalization (Bundel et al. 2016; Jo et al. 2018).

Overall, our results add important context to the existing literature on dopamine’s role in the CeA, particularly how it might influence central pain processing. The inhibitory effects we observed with both D1 and D2 agonists are consistent with the observation of Huang el al. (2022) that stimulating VTA terminals in the CeA has an analgesic effect. Whether the synaptic effects discussed above contribute to pain processing will be an interesting question for future studies. Notably, distinct populations of CeA neurons have been shown to bidirectionally modulate pain behaviors (Wilson et al. 2019) – activating PKCδ^+^ neurons produced mechanical hypersensitivity, while activating SST^+^ neurons reduced sensitivity. Thus, another way dopamine could exert analgesic effects would be to selectively promote the SST^+^ pathway. Indeed, this is precisely the effect observe by Groessl et al. (2018) in the context of freezing behavior, so it is plausible a similar effect could contribute to pain signaling. Improved understanding of these networks could ultimately lead to improved treatment of pain disorders in human patients. It may have particular relevance to conditions with impaired dopamine signaling such as Parkinson disease, in which up to 80% of patients experience pain as a comorbid nonmotor symptom (Andrade et al. 2023; Salabasidou et al. 2024).

## DATA AVAILABILITY

Source data and analysis code is freely available by request to the corresponding author.

## ACKNOWLEDGMENTS

We would like to acknowledge Sherri Vogt, Jakayla Folarin-Hines, and Drew Wolf for invaluable technical assistance with animal colony management.

## DISCLAIMERS

None.

## AUTHOR CONTRIBUTIONS

R.J.H conceived and designed the experiments, performed all experiments and data analysis, prepared figures, and drafted the manuscript. R.G. edited and revised the manuscript. Both authors approved the final version of manuscript.

## REFERENCES

Adke AP, Khan A, Ahn H-S, Becker JJ, Wilson TD, Valdivia S, Sugimura YK, Gonzalez SM, Carrasquillo Y. Cell-Type Specificity of Neuronal Excitability and Morphology in the Central Amygdala. Eneuro ENEURO.0402-20.2020, 2020.

Allen HN, Chaudhry S, Hong VM, Lewter LA, Sinha GP, Carrasquillo Y, Taylor BK, Kolber BJ. A parabrachial-to-amygdala circuit that determines hemispheric lateralization of somatosensory processing. Biol Psychiat, 2022. doi:10.1016/j.biopsych.2022.09.010.

Andrade DC de, Mylius V, Perez-Lloret S, Cury RG, Bannister K, Moisset X, Kubota GT, Finnerup NB, Bouhassira D, Chaudhuri KR, Graven-Nielsen T, Treede R-D. Pain in Parkinson disease: mechanistic substrates, main classification systems, and how to make sense out of them. Pain Publish Ahead of Print, 2023.

Barasi S, Ben-Sreti MM, Clatworthy AL, Duggal KN, Gonzalez JP, Robertson J, Rooney KF, Sewell RDE. Dopamine receptor-mediated spinal antinociception in the normal and haloperidol pretreated rat: effects of sulpiride and SCH 23390. Br J Pharmacol 90: 15–22, 1987.

Bundel DD, Zussy C, Espallergues J, Gerfen CR, Girault J-A, Valjent E. Dopamine D2 receptors gate generalization of conditioned threat responses through mTORC1 signaling in the extended amygdala. Mol Psychiatr 21: 1545–1553, 2016.

Carrasquillo Y, Gereau RW. Hemispheric lateralization of a molecular signal for pain modulation in the amygdala. Mol Pain 4: 24, 2008.

Casey E, Avale ME, Kravitz A, Rubinstein M. Partial Ablation of Postsynaptic Dopamine D2 Receptors in the Central Nucleus of the Amygdala Increases Risk Avoidance in Exploratory Tasks. Eneuro 9: ENEURO.0528-21.2022, 2022.

Casey E, Avale ME, Kravitz A, Rubinstein M. Dopaminergic innervation at the central nucleus of the amygdala reveals distinct topographically segregated regions. Brain Struct Funct 228: 663–675, 2023.

Fernandes MF, Lau D, Sharma S, Fulton S. Anxiety-like behavior in female mice is modulated by STAT3 signaling in midbrain dopamine neurons. Brain, Behav, Immun 95: 391–400, 2021.

Groessl F, Munsch T, Meis S, Griessner J, Kaczanowska J, Pliota P, Kargl D, Badurek S, Kraitsy K, Rassoulpour A, Zuber J, Lessmann V, Haubensak W. Dorsal tegmental dopamine neurons gate associative learning of fear. Nat Neurosci 21: 952–962, 2018.

Guenthner CJ, Miyamichi K, Yang HH, Heller HC, Luo L. Permanent Genetic Access to Transiently Active Neurons via TRAP: Targeted Recombination in Active Populations. Neuron 78: 773–784, 2013.

Hama H, Hioki H, Namiki K, Hoshida T, Kurokawa H, Ishidate F, Kaneko T, Akagi T, Saito T, Saido T, Miyawaki A. ScaleS: an optical clearing palette for biological imaging. Nat Neurosci 18: 1518–1529, 2015.

Huang M, Wang G, Lin Y, Guo Y, Ren X, Shao J, Cao J, Zang W, Li Z. Dopamine receptor D2, but not D1, mediates the reward circuit from the ventral tegmental area to the central amygdala, which is involved in pain relief. Mol Pain 174480692211450, 2022.

Huang S, Zhang Z, Gambeta E, Xu SC, Thomas C, Godfrey N, Chen L, M’Dahoma S, Borgland SL, Zamponi GW. Dopamine Inputs from the Ventral Tegmental Area into the Medial Prefrontal Cortex Modulate Neuropathic Pain-Associated Behaviors in Mice. Cell Reports 31: 107812, 2020.

Ji G, Neugebauer V. Hemispheric lateralization of pain processing by amygdala neurons. J Neurophysiol 102: 2253–64, 2009.

Jin T, Jiang Z, Luan X, Qu Z, Guo F, Gao S, Xu L, Sun X. Exogenous Orexin-A Microinjected Into Central Nucleus of the Amygdala Modulates Feeding and Gastric Motility in Rats. Front Neurosci 14: 274, 2020.

Jo YS, Heymann G, Zweifel LS. Dopamine Neurons Reflect the Uncertainty in Fear Generalization. Neuron 100: 916–925.e3, 2018.

Jokar Z, Khatamsaz S, Alaei H, Shariati M. The electrical stimulation of the central nucleus of the amygdala in combination with dopamine receptor antagonist reduces the acquisition phase of morphine-induced conditioned place preference in male rat. Res Pharm Sci 18: 430–438, 2023.

Kim ES, Lattal KM. Context-Dependent and Context-Independent Effects of D1 Receptor Antagonism in the Basolateral and Central Amygdala during Cocaine Self-Administration. eNeuro 6: ENEURO.0203-19.2019, 2019.

Kim J, Zhang X, Muralidhar S, LeBlanc SA, Tonegawa S. Basolateral to Central Amygdala Neural Circuits for Appetitive Behaviors. Neuron 93: 1464–1479.e5, 2017.

Krishnan B, Genzer KM, Pollandt SW, Liu J, Gallagher JP, Shinnick-Gallagher P. Dopamine-Induced Plasticity, Phospholipase D (PLD) Activity and Cocaine-Cue Behavior Depend on PLD-Linked Metabotropic Glutamate Receptors in Amygdala. Plos One 6: e25639, 2011.

Lançon K, Qu C, Navratilova E, Porreca F, Séguéla P. Decreased dopaminergic inhibition of pyramidal neurons in anterior cingulate cortex maintains chronic neuropathic pain. Cell Reports 37: 109933, 2021.

László K, Péczely L, Géczi F, Kovács A, Zagoracz O, Ollmann T, Kertes E, Kállai V, László B, Berta B, Karádi Z, Lénárd L. The role of D2 dopamine receptors in oxytocin induced place preference and anxiolytic effect. Horm Behav 124: 104777, 2020.

Lee HJ, Youn JM O MJ, Gallagher M, Holland PC. Role of Substantia Nigra-Amygdala Connections in Surprise-Induced Enhancement of Attention. J Neurosci 26: 6077–6081, 2006.

Li J, Chen K, Sheets PL. Topographic organization underlies intrinsic and morphological heterogeneity of central amygdala neurons expressing corticotropin-releasing hormone. J Comp Neurol 530: 2286–2303, 2022.

Lindsay NM, Chen C, Gilam G, Mackey S, Scherrer G. Brain circuits for pain and its treatment. Sci Transl Med 13: eabj7360, 2021.

Liu J, Perez SM, Zhang W, Lodge DJ, Lu X-Y. Selective deletion of the leptin receptor in dopamine neurons produces anxiogenic-like behavior and increases dopaminergic activity in amygdala. Mol Psychiatr 16: 1024, 2011.

Lu J, Cheng Y, Xie X, Woodson K, Bonifacio J, Disney E, Barbee B, Wang X, Zaidi M, Wang J. Whole-Brain Mapping of Direct Inputs to Dopamine D1 and D2 Receptor-Expressing Medium Spiny Neurons in the Posterior Dorsomedial Striatum. Eneuro 8: ENEURO.0348-20.2020, 2021.

Magnusson JE, Fisher K. The involvement of dopamine in nociception: the role of D1 and D2 receptors in the dorsolateral striatum. Brain Res 855: 260–266, 2000.

Mora MP de la, Cárdenas-Cachón L, Vázquez-García M, Crespo-Ramírez M, Jacobsen K, Höistad M, Agnati L, Fuxe K. Anxiolytic effects of intra-amygdaloid injection of the D1 antagonist SCH23390 in the rat. Neurosci Lett 377: 101–105, 2005.

Mora MP de la, Pérez-Carrera D, Crespo-Ramírez M, Tarakanov A, Fuxe K, Borroto-Escuela DO. Signaling in dopamine D2 receptor-oxytocin receptor heterocomplexes and its relevance for the anxiolytic effects of dopamine and oxytocin interactions in the amygdala of the rat. Biochim Biophys Acta (BBA) - Mol Basis Dis 1862: 2075–2085, 2016.

Moscarello JM, Penzo MA. The central nucleus of the amygdala and the construction of defensive modes across the threat-imminence continuum. Nat Neurosci 25: 999–1008, 2022.

Naylor JC, Li Q, Kang-Park M, Wilson WA, Kuhn C, Moore SD. Dopamine attenuates evoked inhibitory synaptic currents in central amygdala neurons. Eur J Neurosci 32: 1836–1842, 2010.

Neugebauer V, Mazzitelli M, Cragg B, Ji G, Navratilova E, Porreca F. Amygdala, neuropeptides, and chronic pain-related affective behaviors. Neuropharmacology 170: 108052, 2020.

Poulin J-F, Caronia G, Hofer C, Cui Q, Helm B, Ramakrishnan C, Chan CS, Dombeck DA, Deisseroth K, Awatramani R. Mapping projections of molecularly defined dopamine neuron subtypes using intersectional genetic approaches. Nat Neurosci 21: 1260–1271, 2018.

Ren W, Centeno MV, Wei X, Wickersham I, Martina M, Apkarian AV, Surmeier DJ. Adaptive alterations in the mesoaccumbal network following peripheral nerve injury. Pain, 2020. doi:10.1097/j.pain.0000000000002092.

Rothman JS, Silver RA. NeuroMatic: An Integrated Open-Source Software Toolkit for Acquisition, Analysis and Simulation of Electrophysiological Data. Front Neuroinform 12: 14, 2018.

Salabasidou E, Binder T, Volkmann J, Kuzkina A, Üçeyler N. Pain in Parkinson disease: a deep phenotyping study. Pain, 2024. doi:10.1097/j.pain.0000000000003173.

Silberman Y, Winder DG. Corticotropin releasing factor and catecholamines enhance glutamatergic neurotransmission in the lateral subdivision of the central amygdala. Neuropharmacology 70: 316–323, 2013.

Smith ES, Fabian P, Rosenthal A, Kaddour-Djebbar A, Lee HJ. The Roles of Central Amygdala D1 and D2 Receptors on Attentional Performance in a Five-Choice Task. Behav Neurosci 129: 564–575, 2015.

Taniguchi W, Nakatsuka T, Miyazaki N, Yamada H, Takeda D, Fujita T, Kumamoto E, Yoshida M. In vivo patch-clamp analysis of dopaminergic antinociceptive actions on substantia gelatinosa neurons in the spinal cord. Pain 152: 95–105, 2011.

Wang Y, Krabbe S, Eddison M, Henry FE, Fleishman G, Lemire AL, Wang L, Korff W, Tillberg PW, Lüthi A, Sternson SM. Multimodal mapping of cell types and projections in the central nucleus of the amygdala. eLife 12: e84262, 2023.

Wilson TD, Valdivia S, Khan A, Ahn H-S, Adke AP, Gonzalez SM, Sugimura YK, Carrasquillo Y. Dual and Opposing Functions of the Central Amygdala in the Modulation of Pain. Cell Reports 29: 332–346.e5, 2019.

Yang T, Yu K, Zhang X, Xiao X, Chen X, Fu Y, Li B. Plastic and stimulus-specific coding of salient events in the central amygdala. Nature 1–10, 2023.

Yates SC, Groeneboom NE, Coello C, Lichtenthaler SF, Kuhn P-H, Demuth H-U, Hartlage-Rübsamen M, Roßner S, Leergaard T, Kreshuk A, Puchades MA, Bjaalie JG. QUINT: Workflow for Quantification and Spatial Analysis of Features in Histological Images From Rodent Brain. Front Neuroinformatics 13: 75, 2019.

Zhang W-H, Zhang J-Y, Holmes A, Pan B-X. Amygdala circuit substrates for stress adaptation and adversity. Biol Psychiat 89: 847–856, 2021.

Zhao Q, Ito T, Soko C, Hori Y, Furuyama T, Hioki H, Konno K, Yamasaki M, Watanabe M, Ohtsuka S, Ono M, Kato N, Yamamoto R. Histochemical characterization of the dorsal raphe-periaqueductal grey dopamine transporter neurons projecting to the extended amygdala. Eneuro, 2022. doi:10.1523/eneuro.0121-22.2022.

Zimmer Z, Fraser K, Grol-Prokopczyk H, Zajacova A. A global study of pain prevalence across 52 countries: examining the role of country-level contextual factors. PAIN 163: 1740–1750, 2022.

